# Defining the Sensitivity Landscape of 74,389 EGFR Variants to Tyrosine Kinase Inhibitors

**DOI:** 10.1101/2021.07.18.452818

**Authors:** Lei An, Shuqing Chen, Guangyao Wu, Chang Liu, Zhenxing Wang, Chunli Wang, Zeyuan Shi, Chenguang Niu, Xiaodong Li, Wenxue Tang, Hongen Xu, Yueqiang Wang

## Abstract

**Background:** Tyrosine kinase inhibitors (TKIs) therapy is a standard treatment for patients with advanced non-small-cell lung carcinoma (NSCLC) when activating epidermal growth factor receptor (*EGFR*) mutations are detected. However, except for the well-studied *EGFR* mutations, most *EGFR* mutations lack treatment regimens.

**Methods:** We constructed two *EGFR* variant libraries containing substitutions, deletions, or insertions using the saturation mutagenesis method. All the variants were located in the *EGFR* mutation hotspot (exons 18–21). The sensitivity of these variants to afatinib, erlotinib, gefitinib, icotinib, and osimertinib was systematically studied by determining their enrichment in massively parallel cytotoxicity assays using an endogenous EGFR-depleted cell line, PC9.

**Results:** A total of 3,914 and 70,475 variants were detected in the constructed *EGFR* Substitution-Deletion (Sub-Del) and exon 20 Insertion (Ins) libraries, accounting for 99.3% and 55.8% of the designed variants, respectively. Of the 3,914 Sub-Del variants, 813 were highly enriched in the reversible TKI (erlotinib, gefitinib, icotinib) cytotoxicity assays and 51 were enriched in the irreversible TKI (afatinib, osimertinib) cytotoxicity assays. For the 70,475 Ins variants, insertions at amino acid positions 770–774 were highly enriched in all the five TKI cytotoxicity assays. Moreover, the top 5% of the enriched insertion variants included a glycine or serine insertion at high frequency.

**Conclusions:** We present a comprehensive reference for the sensitivity of *EGFR* variants to five commonly used TKIs. The approach used here should be applicable to other genes and targeted drugs.

## Background

The incidence of epidermal growth factor receptor (*EGFR*) mutations is reported to be 28.6% among populations with non-small-cell lung carcinoma (NSCLC). East Asian and Southeast Asian populations have a higher incidence (41.3%–57.2%) than the European population (8.0%–20.2%) [1]. Clinical trials have shown that EGFR-tyrosine kinase inhibitor (EGFR-TKI) treatment of patients with NSCLC is superior to chemotherapy in terms of progression-free survival and serious adverse effects [2, 3]. Several EGFR-TKIs (for example, afatinib, erlotinib, gefitinib, and osimertinib) have been approved for first-line treatment of patients with advanced NSCLC having activating *EGFR* mutations [4]. Usually, an *EGFR* mutation testing is performed to check the pattern of mutations in patients, which enables the physicians to determine whether, and which, EGFR-TKI could be used. Statistical analysis of *EGFR* mutation records in the Catalogue of Somatic Mutations in Cancer (COSMIC) database shows that, apart from the secondary mutations (for example, p.Thr790Met, p.Cys797Ser), p.Leu858Arg (41%) and exon 19 deletions (47%) together account for 88% of the mutations in all the records, whereas rare mutations account for about 12% of all mutations [5]. The rare *EGFR* mutations mainly comprise missense variants (>63%) and exon 20 insertions (~17%) [5]. Although rare *EGFR* mutations are reportedly less prevalent, the high incidence of NSCLC has increasingly led to their detection in the clinic [6], making it impossible to ignore them. Unlike for a limited number of high-prevalence *EGFR* mutations, TKI sensitivity for rare mutations and a large number of potential mutations has not been systematically studied.

Several groups have tried to evaluate the clinical relevance of *EGFR* variants on a large scale. One group developed a mixed-all-nominated-mutants-in-one (MANO) method and applied it to 101 nonsynonymous *EGFR* variants [7]. The process of the MANO method is labor intensive, which makes it challenging to evaluate the functions of variants on a large scale, for example, for several thousands of variants. Another group reported screening for activating *EGFR* mutations using a library of 7,216 randomly mutated single-nucleotide variants [8]. This variant library was generated by the error-prone PCR method, which can barely cover all the interested variants.

In view of the lack of economical and efficient methods, it is highly challenging to systematically study the TKI sensitivity of rare *EGFR* mutations on a large scale. Deep mutational scanning (DMS) is a cutting-edge technology that enables the functional analysis of large numbers of variants simultaneously [9, 10]. Usually, a mutant library is first constructed employing synthetic biology or genome editing methods [11, 12], and functional impacts of variants are determined through parallel functional assays. To date, variants of a few clinically actionable genes, such as *BRCA1*, *PPARG*, *TP53*, *PTEN*, *TPMT*, *NUDT15*, *SCN5A*, *CYP2C9*, *CYP2C19*, *CXCR4*, *CCR5*, *ADRB2*, and *MSH2* [12–24], have been extensively studied using the DMS method, highlighting its potential for assessing the TKI sensitivity of rare and potential *EGFR* mutations. In this study, based on the DMS method, we developed massively parallel cytotoxicity assays and systematically determined the sensitivity landscape of the variants to five EGFR-TKIs (Fig. 1).

**Fig. 1.**
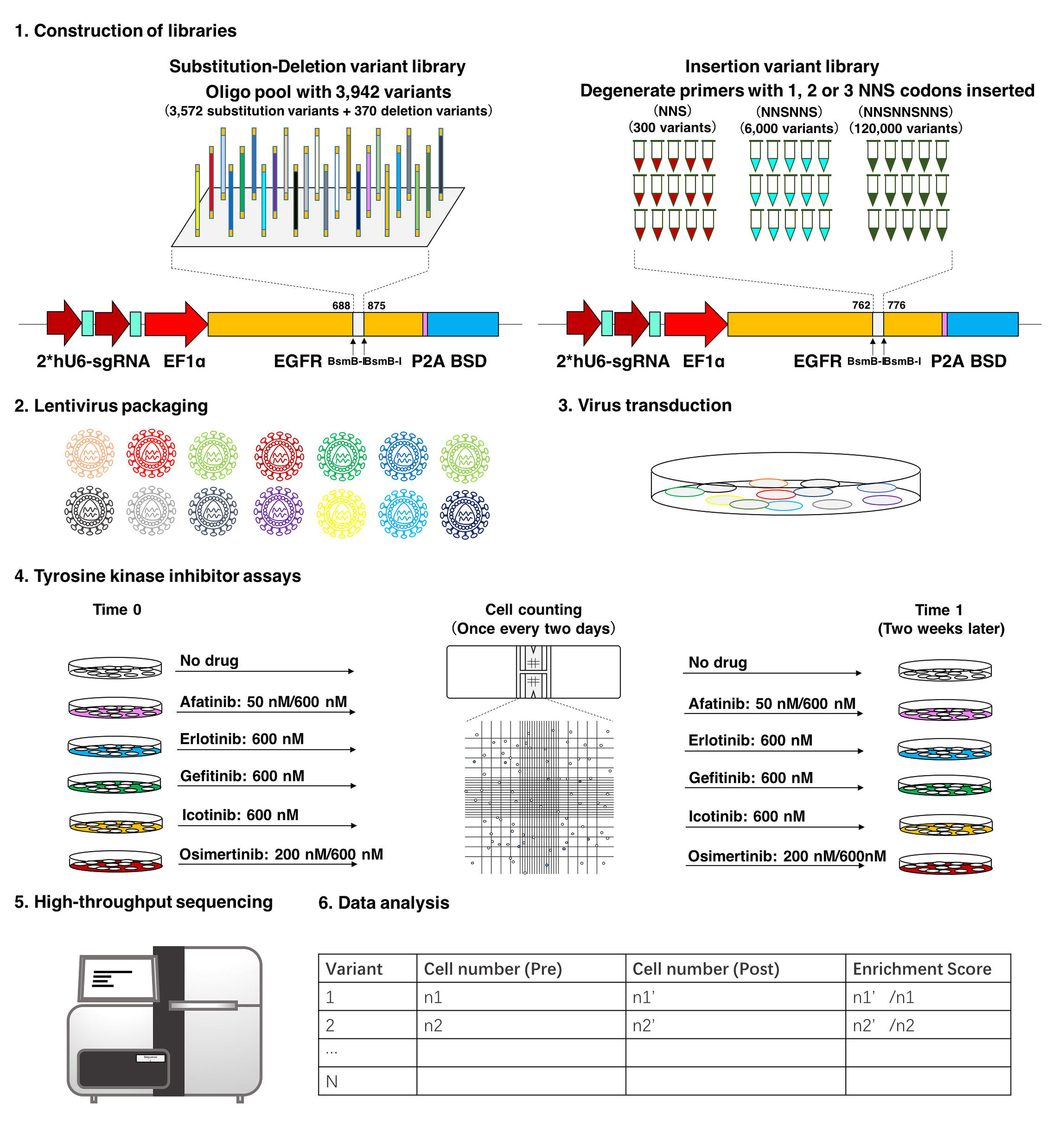
Overview of the experimental design and data analysis. Parallelly synthesized oligonucleotide pools and degenerate primers were separately used to construct *EGFR* substitution-deletion (Sub-Del) and insertion (Ins) variant libraries. Variants in the Sub-Del library were located within *EGFR* exons 18–21 (amino acid positions, 688–875). Variants in the Ins library were located within *EGFR* exon 20 (amino acid positions, 762–776). Each variant had only one designed mutation within a full codon-optimized *EGFR*-coding sequence. Two single-guide RNA (sgRNA) expression cassettes driven by human U6 (hU6) promoter, targeting the endogenous *EGFR* sequence, were located upstream of the *EGFR* variant expression module. A blasticidin S resistance gene (BSD) was fused with the *EGFR* variant cassette through a P2A peptide. The Sub-Del and Ins libraries were separately introduced into Cas9-expressing PC9 cells. The cells were treated with five *EGFR* tyrosine kinase inhibitors (TKIs) or dimethyl sulfoxide (DMSO). The number of cells in each assay was counted manually once, every 2 days. Two weeks later, cells were harvested and subjected to genomic DNA extraction. Mutation regions were amplified by polymerase chain reaction and subjected to next-generation sequencing. Relative drug sensitivity was determined by variant enrichment in cell numbers before and after drug screening.

## Methods

### Cells Lines and Reagents

HEK 293 and A549 cell lines cryopreserved in our laboratory were used. PC9 cells were obtained from Beijing Cancer Hospital and authenticated via short tandem repeat (STR) profiling. All cells were tested to be mycoplasma negative. The genetic background of PC9 cells was further checked by next-generation sequencing using the Illumina TruSight Cancer panel. Cells were maintained according to the instructions prescribed by American Type Culture Collection (ATCC). Unless otherwise noted, all cell culture reagents were purchased from Thermo Fisher Scientific (USA), and all molecular cloning reagents were purchased from New England Biolabs (NEB, USA). Afatinib (S1011), erlotinib (S1023), icotinib (S2922), and osimertinib (S7297) were purchased from Selleck Chemicals and gefitinib (SML1657) was purchased from Sigma-Aldrich. Total EGF Receptor (10001-R021) and anti-EGFR-PE (352904) antibodies were purchased from Sino Biological (China) and BioLegend (USA), respectively.

### CRISPR/Cas9 Single Guide RNA (sgRNA) for Endogenous EGFR Deletion

Ten EGFR targeting sgRNAs were designed using the online tool, CHOPCHOP (https://chopchop.cbu.uib.no/). The sgRNAs cloning were performed according to the reported protocol [25]. CRISPR/Cas9 lentiviral particles were separately produced and then introduced into A549 cells. The gene editing efficiency of the sgRNAs was determined using fluorescent-activated cells sorting (FACS) analysis and immunoblotting.

### Construction of Saturation Mutagenesis Libraries

*EGFR* saturation mutagenesis libraries were constructed based on the Mutagenesis by Integrated TilEs (MITE) method reported previously [11], with necessary modifications. The complete Substitution-Deletion (Sub-Del) Library was obtained by mixing six sub-libraries at equal mass ratios. The complete Insertion (Ins) Library was obtained by mixing three sub-libraries (Ins-1, Ins-2, Ins-3) in the mass ratio of 1:5:5. The completeness and uniformity of the variant libraries were verified through next-generation sequencing using a HiSeq sequencer (Illumina).

### Lentivirus Production

Lentivirus was produced and titered as reported previously [15], with minor modifications. For lentivirus production, five million 293T cells were pre-seeded on 10 cm dishes, the first day evening. Cells were cotransfected with the variant library plasmid and two helper plasmids (psPAX2 and pMD2.G) at a molar ratio of 1:1:1 using a cationic polymer transfection reagent, EZ Trans (LIFE iLab, China), according to the manufacturer’s instructions. The medium was replaced 24 h later and the virus particles were collected and filtered through 0.45 μm filters (Millipore), 24 or 48 h later.

### TKI Cytotoxicity Screening

PC9 cells were transduced with a Cas9 expression lentivirus and selected on puromycin for 2 weeks. PC9 cells stably expressing Cas9 (PC9-Cas9) were cultured in large quantities and seeded in twenty 15 cm dishes (~15 million cells per dish), the first day evening. Subsequently, these cells were transduced with a pooled virus of the Sub-Del/Ins library (10 dishes for each library) at a multiplicity of infection of ~0.25. Transduced cells were selected on a medium with puromycin and blasticidin for 2 weeks. For the TKI sensitivity screening, cells transduced with the Sub-Del/Ins library were separately seeded in eight 15 cm dishes (15 million cells per dish) and treated with different TKIs or dimethyl sulfoxide (DMSO): two for afatinib (50 nM/600 nM), one for erlotinib (600 nM), one for gefitinib (600 nM), one for icotinib (600 nM), two for osimertinib (200 nM/600 nM), and one for DMSO (control). The remaining cells (>5 million cells for each library) were collected as the time 0 samples. Cells were trypsinized, manually counted, and reseeded every 2 days for 14 days. Dishes of proper size were used each time to ensure that the cell density was maintained within a proper range. The TKI assays were independently repeated once (Additional file 1: Figure S1). After 14 days of TKI treatment (time 1), cells were harvested for genomic DNA (gDNA) extraction.

### Amplification and Sequencing of Mutational Regions

The first-round PCR was performed to amplify a ~1.1 kb DNA fragment covering the entire *EGFR* mutational region from the gDNA. Multiple 50 μL PCRs were performed for each sample. PCR products from each sample were separately pooled and purified with a Universal DNA Purification Kit (Tiangen, China). In the second-round PCR, the Sub-Del library mutational region was amplified with corresponding tagged primers (Additional file 2: Table S1) using the 1.1 kb fragments as templates. In contrast, the Ins library mutational region was amplified from the corresponding templates. All PCRs were performed using the Q5 High-Fidelity DNA Polymerase (NEB). The amplicons were purified using a Universal DNA Purification Kit. Next-generation sequencing libraries were constructed according to the manufacturer’s protocol (VAHTS, China). All samples were sequenced on the HiSeq platform (Illumina) using a 2×150 paired-end configuration.

### Data Processing

Specifically, raw overlapping paired reads (read1 and read2) were merged into single reads using the bbmerge tool with default parameters [26]. For Sub-Del libraries, each read was assigned to one reference sequence using the USEARCH tool with parameters of -usearch_global -strand both -id 0.95 [27]. For Ins libraries, each read was assigned to one reference sequence with the same USEARCH tool but with a different value of id parameter (0.90). The successfully assigned reads in Sub-Del and Ins libraries were translated into AA and were subjected to diamond blastp analysis against the reference protein sequences to call variants [28]. The frequency of each mutation was calculated as the ratio of the variant count to the count of valid variants. Thereafter, the frequency of each variant was weighted with the ratio of cell count change relative to time 0. To assess the enrichment or depletion of each variant, we calculated the log2-fold change in variant frequency relative to the time 0 samples. Heatmap and violin plots were created using the R pheatmap and ggplot2 packages, respectively.

## Results

### Construction of 74,389 EGFR Variants

*EGFR* pathogenic variants, including substitutions, deletions, and exon 20 insertions [29], are mainly located in exons 18–21 (residues 688–875). To cover all the major types of mutations, two *EGFR* libraries were constructed using the MITE method [11]. Parallel synthesized oligonucleotides were used to construct a saturation *EGFR* Sub-Del library containing 3,572 (theoretical number) single AA substitution (Sub) variants, 188 (theoretical number) single AA deletion (Del-1) variants, and 182 (theoretical number, for cloning reason, six deletion variants were not covered) two consecutive AA deletion (Del-2) variants. All Sub-Del library variants were located at AA positions 688–875. Similarly, multiple degenerate primers were used to construct a saturation *EGFR* Ins library containing 126,300 (theoretical number) in-frame insertion variants. All Ins library variants were located in the EGFR insertion hotspot (AA position, 762–776) and consisted of one, two, or three tandem NNS codons (N = adenine, cytosine, guanine, or thymine; S = cytosine or guanine) inserted between adjacent AAs, with each variant containing one insertion in each of the AA positions. The Ins library was composed of three subtype libraries: Ins-1 library (300 variants, each with a single-AA inserted), Ins-2 library (6,000 variants, each with a double-AA inserted), and Ins-3 library (120,000 variants, each with a triple-AA inserted). Variants of Sub-Del and Ins libraries were cloned into lentiviral vectors, and the uniformity and completeness of each library were determined by high-throughput sequencing of the plasmids. The Sub-Del library showed decent uniformity (Additional file 3: Figure S2a), and 99.3% of the designed variants were detected more than 20 times (>20 reads) (Additional file 4: Figure S3a). Similarly, variants of the Ins library also showed high uniformity (Additional file 3: Figure S2b). Because the Ins library was derived from mixing the Ins-1, Ins-2, and Ins-3 libraries at a mass ratio of 1:5:5, the ratio of the average proportion of each variant from Ins-1, Ins-2, and Ins-3 libraries was 80:20:1. Not surprisingly, most of the designed variants in the Ins-1 (99.7%) and Ins-2 (93.6%) libraries were detected (>20 reads for each variant). However, only 53.8% of the designed Ins-3 library variants were detected (>20 reads for each variant) (Additional file 4: Figure S3b). Overall, the coverage and uniformity of Sub-Del and Ins libraries were as expected, and the total number of detected variants was 74,389.

### Deletion of Endogenous EGFR in the PC-9 Cell Model

The human lung adenocarcinoma cell line, PC9 (contains *EGFR* exon 19 deletions), was chosen as the model. The cell line identity was confirmed by STR profiling and genomic mutations were further reviewed through next-generation sequencing. A Glu746–Ala750 deletion was confirmed in *EGFR* exon 19 and a p.Arg248Gln mutation was detected in *TP53*. The p.Arg248Gln variant is a gain-of-function mutation that promotes tumorigenesis [30]. These results indicate that the PC9 cell line is a genetically clear cell model for evaluating the sensitivity of *EGFR* variants to TKIs.

To eliminate the endogenous EGFR, we separately introduced Cas9 and the sgRNA-*EGFR* variant (codon-optimized) into PC9 cells using two lentiviral vectors. Firstly, Cas9 was introduced into PC9 cells to obtain the PC9-Cas9 cell line, stably expressing Cas9. Subsequently, the sgRNA-*EGFR* variant was introduced into the PC9-Cas9 cells. The endogenous EGFR was knocked out using the CRISPR/Cas9 system and replaced with the exogenous EGFR variant in approximately 10 days. Because direct knockout of EGFR may seriously affect the viability of PC9 cells [31], we tested the CRISPR/Cas9 gene-editing efficiency in A549 cells (with a wildtype *EGFR* and mutated *KRAS* background). Considering that *KRAS* locates downstream of the *EGFR* signaling pathway, *EGFR* can be knocked out without seriously affecting the viability of A549 cells. FACS and western blotting analyses confirmed that the endogenous EGFR had been efficiently knocked out by CRISPR/Cas9 (Additional file 5-7: Figure S4, S5; Table S2). We predicted that these validated CRISPR/Cas9 sgRNAs would also efficiently knockout the endogenous EGFR in PC9-Cas9 cells.

### EGFR-TKI Cytotoxicity Screening

To systematically evaluate the sensitivity of *EGFR* variants to different TKIs, we separately introduced Sub-Del and Ins variants into the PC9-Cas9 cells and treated them with TKIs. Expression of exogenous EGFR variants was checked through FACS analysis and found to be slightly higher than that of EGFR in PC9 cells (Additional file 8: Figure S6). The concentrations of EGFR-TKIs were set mainly referring to the clinical plasma concentration of each TKI (Additional file 9: Table S3). Specifically, the concentrations of reversible TKIs (erlotinib, gefitinib, and icotinib) were set to 600 nM, whereas those of irreversible TKIs were set at two levels. The afatinib clinical plasma concentration is much lower than 600 nM [32]; we designed two assays for 50 and 600 nM afatinib. Because incubation with 600 nM osimertinib showed strong cytotoxicity to cells with Sub-Del variants *in vitro*, we designed two assays for 200 and 600 nM osimertinib. The TKI treatments lasted 2 weeks and cells were counted every 2 days. The number of cells containing the Sub-Del variants in the erlotinib, gefitinib, icotinib, afatinib (50 nM), and osimertinib (200 nM) assays decreased substantially at days 5 and 7, reached the minimum at day 9, and then began to increase. The cell numbers in the afatinib (600 nM) and osimertinib (600 nM) assays reached the minimum at days 11 and 15, respectively (Fig. 2a). For cells containing the Ins variants, the minimum cell number in the afatinib (600 nM) and osimertinib (600 nM) assays was reached at days 9 and 13, respectively, whereas the cell number in other TKI assays was not decreased during 2-week of TKI treatment (Fig. 2b).

**Fig. 2.**
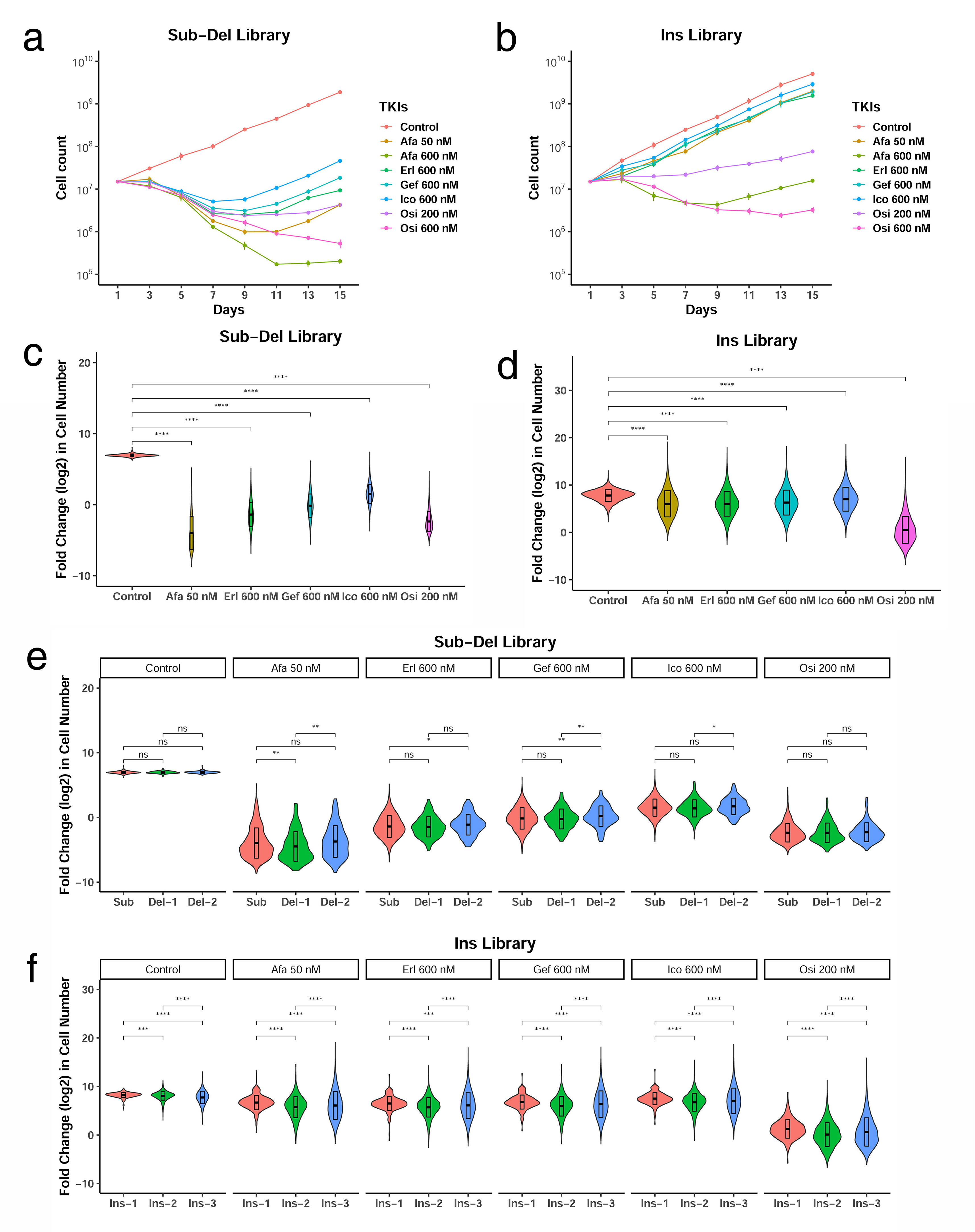
*EGFR*-tyrosine kinase inhibitors (EGFR-TKIs) showed different inhibition on Sub-Del and Ins library variants. Cell growth kinetics (log10) of the Sub-Del (**a**) and Ins (**b**) libraries treated with five TKIs and dimethyl sulfoxide (DMSO) (mechanical replicates, n = 4; error bars indicate the standard deviation). Violin plots denoting enrichment score (log2) distributions of variants in the Sub-Del (**c**) and Ins (**d**) libraries treated with the five TKIs or DMSO. **e** The enrichment scores (log2) of the three subtypes of variants (Sub: single amino acid (AA) substitution; Del-1: single AA deletion; and Del-2: dual AA deletion) in the Sub-Del library are separately displayed by treatment type in the violin plot. **f** The enrichment scores (log2) of the three subtypes of variants (Ins-1: single AA insertion; Ins-2: dual AA insertion; and Ins-3: triple AA insertion) in the Ins library are separately displayed by treatment type in the violin plot. Student’s *t*-test, ns: P > 0.05, *: P ≤ 0.05, **: P ≤ 0.01, ***: P ≤ 0.001, ****: P ≤ 0.0001.

### Osimertinib Inhibited Almost all Types of Variants

To investigate the enrichment of variants after incubation with TKIs, cells from each TKI assay were harvested for gDNA extraction, mutation region amplification, and high-throughput sequencing. The relative drug sensitivity for each variant was defined as the fold change of a variant’s theoretical cell number post-TKI screening (time 1) to that at the early time point (time 0). The theoretical cell number for each variant was calculated using the read proportion of a variant among the total valid reads multiplied by the final cell number in each assay. Compared with reversible TKIs (erlotinib, gefitinib, and icotinib), irreversible TKIs (afatinib 50 nM and osimertinib 200 nM) showed stronger inhibition for Sub-Del variants (enrichment score mean: afatinib, 0.331; erlotinib, 0.793; gefitinib, 1.73; icotinib, 4.44; osimertinib, 0.388; false discovery rate (FDR)-adjusted P values <0.0001) (Fig. 2c). For the Ins variants, osimertinib (200 nM) had a greater inhibition than the other TKIs (enrichment score mean: afatinib, 113; erlotinib, 89.3; gefitinib, 111; icotinib, 176; osimertinib, 4.46; FDR-adjusted P values <0.0001) (Fig. 2d). Overall, these results indicated that the Ins library variants were more resistant to TKI treatment than the Sub-Del library variants, and osimertinib (200 nM) showed stronger inhibition for variants in both the libraries.

To investigate whether distinct *EGFR* variant subtypes present different sensitivity against the same EGFR-TKI, we separately analyzed relative drug sensitivity of the Sub, Del-1, and Del-2 variants in the Sub-Del library and the Ins-1, Ins-2, and Ins-3 variants in the Ins library. No significant difference in the inhibition of Sub, Del-1, and Del-2 variants to any of the five TKIs was observed (Fig. 2e, FDR-adjusted P values >0.05). In contrast, the sensitivity of Ins-1, Ins-2, and Ins-3 variants to each of the five TKIs was significantly different (Fig. 2f, FDR-adjusted P values < 0.05). However, the difference in enrichment scores (log2) cannot be entirely attributed to TKI treatment because differences were also observed for Ins-1, Ins-2, and Ins-3 variants in TKI-free assay (control).

### Cytotoxicity Screening Results were Consistent with Known Clinical Annotations

Next, we evaluated the consistency of TKI screening results with known clinical annotations. For this, we compiled a list of *EGFR* mutations from the Clinical Interpretation of Variants in Cancer (CIViC) knowledgebase and compared the clinical annotations (resistant or sensitive) of mutations with the enrichment scores (log2) obtained in cytotoxicity assays by plotting those mutations on a scale bar comprising all the results (Additional file 10: Figure S7). TKI-sensitive variants (such as p.Leu858Arg and p.Gly719Ala/Ser) were located in the middle or lower half of the bar, whereas TKI-resistant variants (such as p.Thr790Met) were located in the middle or upper half of the bar (Additional file 10: Figure 7a). Similarly, the annotated Ins variants were distributed at expected positions in the bars (Additional file 10: Figure 7b). However, some annotated variants, such as p.Cys797Ser (0.53) and p.Ser768Ile (1.19), had contradictory enrichment scores in the osimertinib assay. These discrepancies could be attributed to the presence of additional mutations. For example, *EGFR* mutation combinations in *cis* (on the same allele) or *trans* (on different alleles) might lead to altered drug response outcomes compared with those for independent mutations [33–36]. Indeed, p.Cys797Ser has been detected as a secondary mutation coexisting with p.Thr790Met in *cis* and results in resistance to osimertinib [33–35]. In addition, another variant, p.Gly724Ser, has been associated with increased sensitivity to osimertinib when combined in *trans* with p.Leu858Arg than with exon 19 deletions [36]. In general, our TKI sensitivity evaluation results are consistent with the known annotation.

### Cytotoxicity Screening Identifies Potentially Drug Resistant Variants

The variant enrichment scores (log2) in each TKI assay for each variant were presented as heatmaps (**Fig. 3, 4**, Additional file 11, 12: Figure S8, S9). For Sub-Del variants, 280 (7.2%) variants in afatinib assay (50 nM), 841 (21.5%) variants in erlotinib assay, 1,802 (46.2%) variants in gefitinib assay, 3,440 (88.0%) variants in icotinib assay, and 271 (7.2%) variants in osimertinib assay (200 nM) of all variants had a positive enrichment score (log2), indicating that the number of those variants were increased after 2 weeks of TKI incubation. We divided all substitution variants into two categories, the complex and simple variants. The complex variants can only be achieved through two or three point mutations in the corresponding codon whereas the simple variants can be obtained through one point mutation. Under normal circumstances, the probability of multiple independent point mutations occurring in the same codon is relatively low, and therefore, simple variants are more likely to be encountered in the clinic. Enriched variants from all three reversible TKI assays have 813 variants (complex variants = 571, simple variants = 242) in common (Fig. 5 a, c, e). The variants enriched in the two irreversible TKI assays have 51 variants (complex variants = 36, simple variants =15) in common (Fig. 5 b, d, f). Overall, in our systematic cytotoxicity screening, we found 813 and 51 substitution variants that potentially mediate resistance to *EGFR* reversible and irreversible TKIs, respectively.

**Fig. 3.**
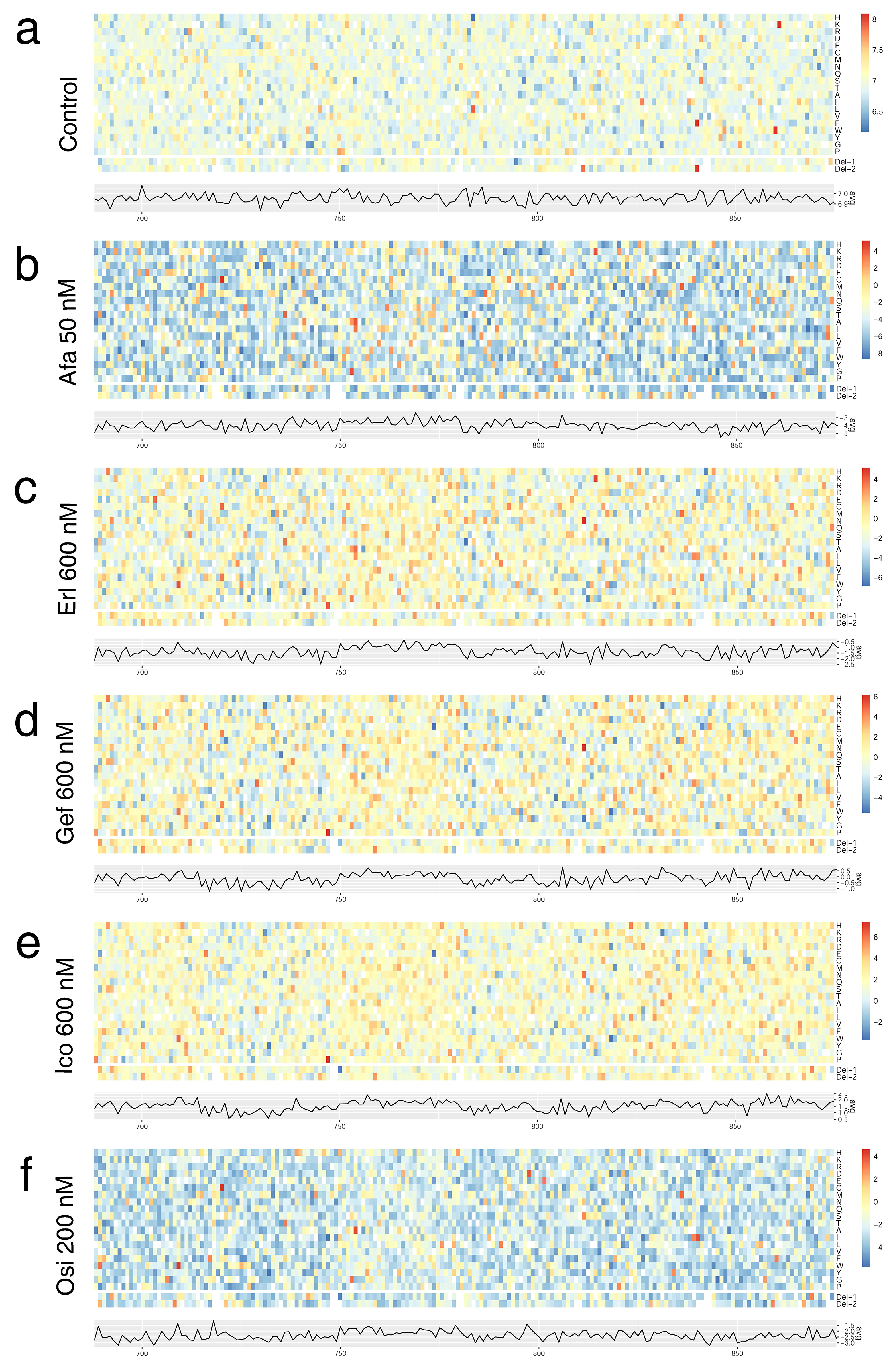
Heatmaps depicting enrichment of *EGFR* Sub-Del library variants on the cytotoxicity of five tyrosine kinase inhibitors (TKIs). The enrichment scores are shown as the ratio (log2) of the cell number post-TKI treatment to that pre-TKI treatment. The top 20 rows represent all 20 possible amino acid (AA) substitutions, and the two rows below represent one or two AA deletion mutations (Sub: single AA substitution; Del-1: single AA deletion; and Del-2: dual AA deletion). Each column indicates one AA residue in EGFR (688–875). Blue to red represents high to low abundance of variants, respectively. Variants that were not covered are colored in white. Codon-level average enrichment scores are plotted below each heatmap. Cells were treated with dimethyl sulfoxide (DMSO) (**a**), 50 nM afatinib (**b**), 600 nM erlotinib (**c**), 600 nM gefitinib (**d**), 600 nM icotinib (**e**), or 200 nM osimertinib (**f**).

**Fig. 4.**
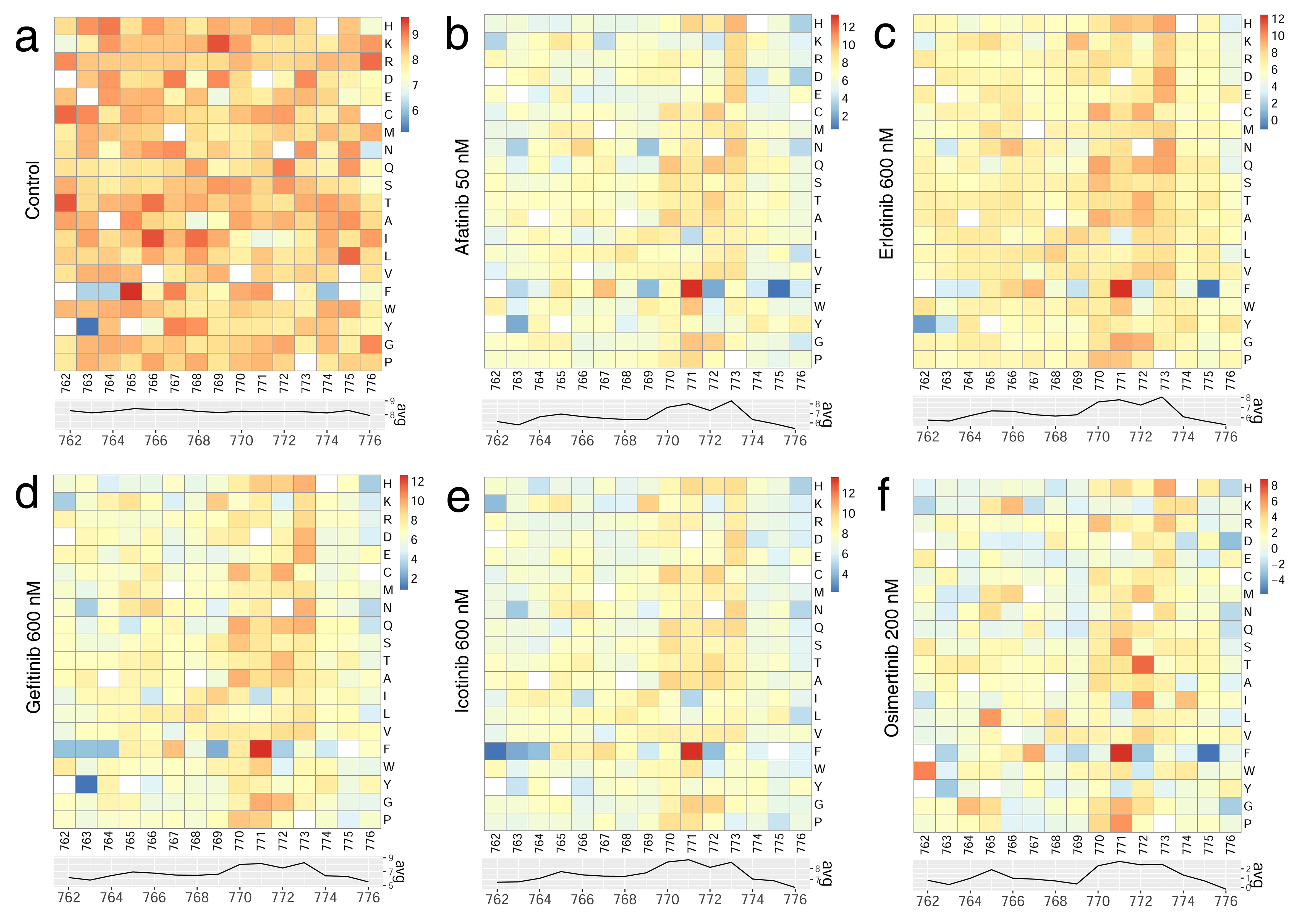
Heatmaps depicting the enrichment results for *EGFR* Ins-1 library variants on the cytotoxicity of five tyrosine kinase inhibitors (TKIs). The enrichment scores are shown as the ratio (log2) of the cell number post-TKI treatment to that pre-TKI treatment. The 20 rows represent all 20 possible single AA insertion mutations. Each column indicates one AA residue in EGFR (762–776). Blue to red represents high to low abundance of variants, respectively. Variants that were not covered are colored in white. Codon-level average enrichment scores are plotted below each heatmap. Cells were treated with dimethyl sulfoxide (DMSO) (**a**), 50 nM afatinib (**b**), 600 nM erlotinib (**c**), 600 nM gefitinib (**d**), 600 nM icotinib (**e**), or 200 nM osimertinib (**f**).

**Fig. 5.**
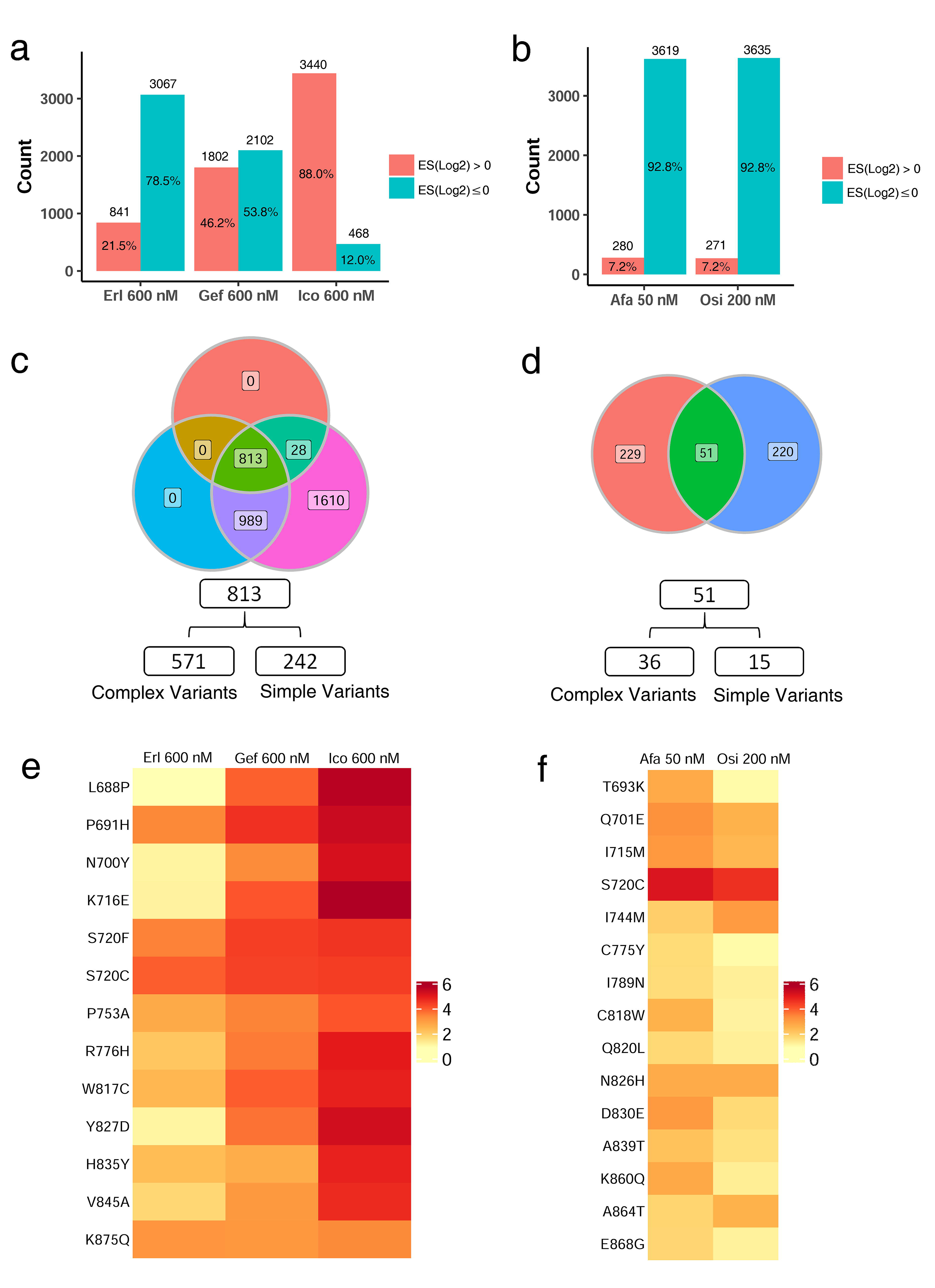
Potentially drug-resistant substitutions identified in the cytotoxicity screening. Number and proportion of enriched (red bar) and depleted (green bar) variants in the reversible (**a**) and irreversible (**b**) tyrosine kinase inhibitor (TKI) assays. **c** The enriched variants in the three reversible TKI assays have an intersection of 813 variants (571 complex and 242 simple variants). **d** The enriched variants in the two irreversible TKI assays have an intersection of 51 variants (36 complex and 15 simple variants). **e** The enrichment score (log2) of the top 5% of the selected simple variants in “**c**” is presented as a heatmap. **f** The enrichment score (log2) of all the 15 selected simple variants in “**d**” was presented as a heatmap. Afa, Erl, Gef, Ico, Osi, and ES are abbreviations for afatinib, erlotinib, gefitinib, icotinib, osimertinib, and enrichment score, respectively.

### The Top 5% Enriched Insertions Showed Insertional Position and AA Type Preference

For Ins variants, the proportion of variants with positive enrichment scores (log2) was much larger than of their Sub-Del counterparts: 95% for afatinib (50 nM) assay, 93% for erlotinib assay, 94% for gefitinib assay, 97% for icotinib assay, and 57% for osimertinib (200 nM) assay. To study whether the insertional position and the AA type affect the sensitivity of exon 20 insertions to different TKIs, we selected the top 5% enriched insertions from each TKI assay and determined the cumulative frequency of the inserted AA at each AA position. The top enriched exon 20 insertions shared many common features (Fig. 6 a-f): (1) the highly enriched insertion variants showed preference for insertion at AA positions 770–774; (2) the highly enriched insertion variants showed preference of glycine and serine. To evaluate the sensitivity of the clinically detected exon 20 insertions to TKIs, we determined the cumulative frequency of insertional AA in variants from the Chinese population [37] and COSMIC database (Fig. 6 g-i). The detected rare exon 20 insertions were mainly concentrated within AA positions 770–774. Our results explain the resistance of these clinically identified exon 20 insertions to TKIs. Together, these results reveal that insertion in the *EGFR* AA positions 770–774 might induce a stronger resistance to TKIs, especially when the insertions contain glycine or serine.

**Fig. 6.**
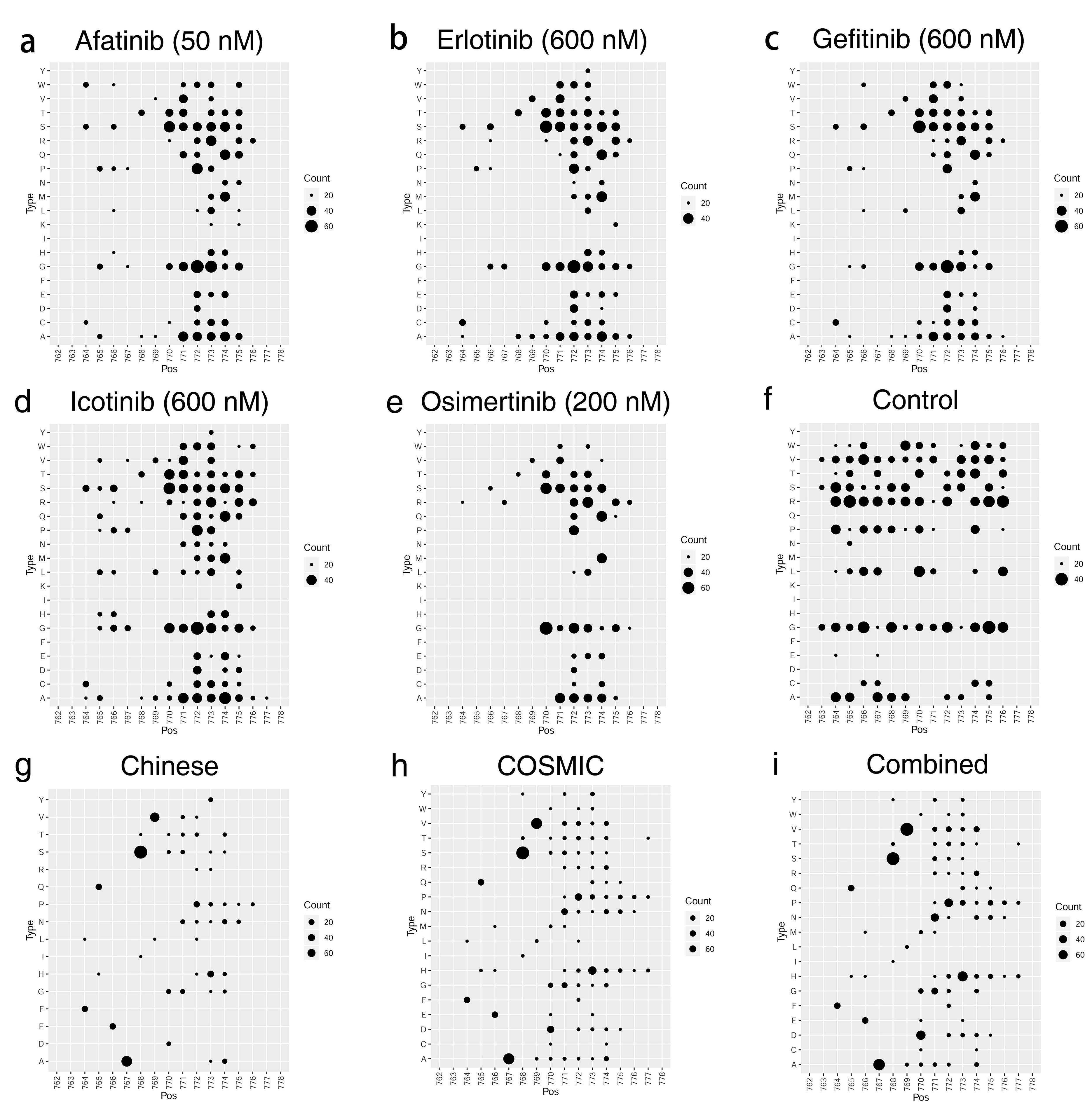
Variants in *EGFR* exon 20 insertional hotspot induce stronger tyrosine kinase inhibitor (TKI) resistance. **a-f** The inserted amino acids (AA) in the top 5% enriched exon 20 insertions were counted and the cumulative frequency numbers were plotted as black dots at the corresponding AA positions. Dots with cumulative frequency less than 15 were filtered. **a-e** show summary results for afatinib (50 nM), erlotinib, gefitinib, icotinib, and osimertinib (200 nM) assays, respectively. **f** shows the summary result for untreated control. **g-i** The inserted AA in exon 20 insertions from the Chinese population [37] and COSMIC database were counted and the cumulative frequency numbers were plotted as black dots at the corresponding AA positions. All the dots are shown, regardless of the cumulative frequency numbers. **g** shows the summary result for the Chinese population. **h** shows the summary result for the COSMIC database. **i** shows combined results of “**g**” and “**h**”. The X-axis shows AA positions.

## Discussion

In the post-genome era, a massive number of variants of unknown significance (VUS) has been identified by high-throughput sequencing; however, it is still a big challenge for functional interpretation of VUS in the clinical context. To study the TKI sensitivity of rare *EGFR* mutations, we designed cytotoxicity assays and systematically evaluated the sensitivity of 74,389 *EGFR* variants to five commonly used EGFR-TKIs. Afatinib and osimertinib showed intense and relatively persistent inhibition of all types of *EGFR* variants (substitutions, deletions, and exon 20 insertions) among the five tested EGFR-TKIs. Considering the tolerance to the treatment, the clinical plasma drug concentration of afatinib cannot reach 600 nM [38], whereas for osimertinib, it can be more than 600 nM [39]. Accordingly, we speculate that otherwise for additional evidence, patients with NSCLC having rare *EGFR* mutations (including exon 20 insertions) are more likely to benefit from osimertinib than from the other four TKIs. Compared with Sub-Del mutations, *EGFR* exon 20 insertion mutations were more difficult to deal with. A recent clinical study showed that patients with *EGFR* exon 20 insertions could benefit from high-dose (160 mg daily) osimertinib treatment [40]. With the combined use of synthetic biology and CRISPR/Cas9 methods, we can economically generate a large number of cell models with designed mutations of a specific gene. These cell models can partially simulate clinical samples and enable us to obtain rapid screening results through appropriate functional assays.

The systematic cytotoxicity screening of Sub-Del variants has led us to identify several potentially drug-resistant substitution variants. We were curious whether these variants occurred with equal frequency in the clinic. Further analysis revealed that the substitution variants could be divided into complex and simple variants. The majority of *EGFR* substitution mutations detected in the clinic are simple variants and most of the complex variants are rarely detected. The difficulty in the mutation process partially explains why the highly enriched complex variants are rarely detected in the clinic [15]. To the best of our knowledge, we have designated substitution variants into complex and simple variants for the first time. More efforts should be invested on the functional interpretation of the simple variants in the clinical context.

The systematic screening revealed that *EGFR* exon 20 insertions at AA positions 770–774 would induce stronger TKI resistance, especially when the insertions contain glycine or serine. Clinically, excluding the high prevalence exon 20 insertions of p.Ala767_Val769dup and p.Ser768_Asp770dup, a large number of low prevalence exon 20 insertions are mostly concentrated at AA positions 770–774 [37, 41]. According to the theory of Vyse *et al*., *EGFR* exon 20 insertions in the loop region (AA positions 767–775) have altered three-dimensional structures and will stabilize the mutated EGFR in the active state even without ligand binding [41]. Our systematic screening results provided important experimental evidence supporting this theory.

The cytotoxicity assay-based method for functional interpretation of variants used in this study can also be applied to other genes and targeted drugs. At least 52 small molecule kinase inhibitors have been approved by US food and drug administration for targeted therapy [42]. An extensive functional interpretation of variants will enable us to clarify the targeted inhibitors applicable for the corresponding mutations. Moreover, high throughput sequencing of circulating tumor DNA allows us to forecast the potential drug-resistance events [43–46]. This will dramatically promote the development of precision medicine. Moreover, for newly developed small molecule targeted inhibitors, a pre-drug sensitivity screening at the laboratory stage will help forecast the suitability of drug candidates for patients with specific mutations, thereby drastically reducing cost incurred at clinical trial stages.

Despite the significance of this work, there are several important caveats: (1) We only used PC9 cells for cytotoxicity screening. The genetic background of this cell line might have had an unwanted influence on the functional interpretation of variants [12, 19]. Use of multiple NSCLC cell lines for parallel functional screening can significantly reduce such interference; (2) The expression of the exogenous *EGFR* variants was higher than that of endogenous *EGFR* in PC9 cells, which may lead to an overestimation of the TKI resistance of specific variants; (3) The endogenous *EGFR* of PC9 cells was deleted by CRISPR/Cas9. In theory, each of the engineered PC9 cells has one copy of the exogenous *EGFR* variant. However, several studies have demonstrated that *EGFR* variant combinations can significantly influence sensitivity to a specific TKI [33–36, 47]. Further studies are needed to uncover the TKI inhibition against combinations of *EGFR* variants.

## Conclusions

In conclusion, systematic cytotoxicity screening revealed that patients with NSCLC having rare *EGFR* mutations are most likely to benefit from osimertinib treatment. We expect variants of other genes to be classified to assess their sensitivity to corresponding targeted inhibitors. All these efforts will greatly promote the development of cancer precision medicine.

## Abbreviations

AA: Amino acid
Afa: Afatinib
ATCC: American Type Culture Collection
CIViC: Clinical Interpretation of Variants in Cancer
COSMIC: Catalogue of Somatic Mutations in Cancer
DMS: Deep mutational scanning
DMSO: Dimethyl sulfoxide
EGFR: Epidermal growth factor receptor
EGFR-TKI: EGFR-tyrosine kinase inhibitor
Erl: Erlotinib
FACS: Fluorescent-activated cells sorting
FDR: False discovery rate
gDNA: Genomic DNA
Gef: Gefitinib
Ico: Icotinib
Ins: Insertion
MANO: Mixed-all-nominated-mutants-in-one
MITE: Mutagenesis by Integrated TilEs
NEB: New England Biolabs
NSCLC: Non-small-cell lung carcinoma
Osi: Osimertinib
sgRNA: Single guide RNA
STR: Short tandem repeat
Sub-Del: Substitution-Deletion
TKI: Tyrosine kinase inhibitor
VUS: Variants of unknown significance

## Declarations

### Ethics approval and consent to participate

Not applicable.

### Consent for publication

Not applicable.

### Availability of data and materials

The datasets are available from the corresponding authors on reasonable request.

### Authors’ Contributions

YW, LA, HX and SC conceived the work and designed the experiments; YW, LA, SC, GW, CL, ZW, CW, and ZS performed the experiments; LA, HX, CN, XL, and WT provided funding and expertise support; HX, SC, and YW analyzed the data; YW wrote the original manuscript; YW, LA, HX and SC contributed to the manuscript revision. All authors read and approved the final manuscript.

## Acknowledgments

We thank Dr. Kang Bin from Cancer Hospital of Peking University for PC9 cell line and Luo Yuqi for experimental assistance. We thank Dr. Ren Yanfang of the University of Rochester for his comments on the manuscript. The data analysis was supported by the Supercomputing Center at Zhengzhou University (Zhengzhou).

